# Copy number aberrations drive kinase re-wiring leading to genetic vulnerabilities in cancer

**DOI:** 10.1101/515932

**Authors:** Danish Memon, Michael B. Gill, Eva Papachristou, David Ochoa, Clive D’Santos, Martin L. Miller, Pedro Beltrao

## Abstract

Somatic DNA copy number variations (CNVs) are prevalent in cancer and can drive cancer progression albeit with often uncharacterized roles in altering cell signaling states. Here, we integrated genomic and proteomic data for 5598 tumor samples to identify CNVs leading to aberrant signal transduction. The resulting associations recapitulated known kinase-substrate relationships and further network analysis prioritized likely driver genes. A total of 44 robust pan-cancer gene-phosphosite associations were replicated in cell line samples. Of these, ARHGEF17, a predicted regulator of hippo-signaling, was further studied through (phospho)proteomics analysis where ARHGEF17 knockdown cells showed dys-regulation of hippo- and p38 signaling as well as immune related pathways. Using, RNAi, CRISPR and drug screening data we find evidence of kinase addiction in cancer cell lines identifying inhibitors for targeting of ‘kinase-dependent’ cell lines. We propose copy number status of genes as useful predictors of differential impact of kinase inhibition, a strategy that may be of use in the future for anticancer therapies.

## Introduction

Kinases are druggable proteins that are key targets for cancer treatment as they are highly prone to acquire somatic mutations and act as oncogenes (Gross et al., 2015). Several studies have proposed that cancer cells can become dependent or addicted to changes in kinase signaling (Sawyers, 2004). However, the challenge remains to identify the genomic context in which specific kinase inhibitors are more likely to be effective. Cancer genomes harbour a large number and variety of mutations including somatic point mutations and copy number aberrations. These mutations may have many direct and indirect effects which are likely to rewire signaling pathways giving cancer cells a growth advantage. Several studies have reported evidence of somatic mutations that affect kinase activity (Miller et al., 2015; Olow et al., 2016; Reimand and Bader, 2013). However, the impact of copy number aberrations on kinase signaling activity are often unknown besides a some well characterized cases of CNV in signaling genes such as PTEN (Chalhoub and Baker, 2009) and HER2 (Moasser, 2007). Several tumor types including breast (Curtis et al., 2012) and ovarian cancer (Macintyre et al., 2018) show large-scale genomic aberrations and have been known to contribute to cancer development and progression. Identifying downstream effects of copy number changes is complicated by the fact that they encompass large segments of genomes with many genes and therefore it is difficult to identify the likely ‘driver’ gene within the CNV region.

Large scale phosphorylation measurements of tumor samples have relied primarily on Reverse Phase Protein Arrays (RPPA), that consists of a (phospho-)protein antibody microarrays used to estimate protein abundances in a high-throughput manner. The TCGA RPPA platform (TCPA) currently has estimates for around 150 proteins and 50 phospho-proteins for nearly 10,000 tumor samples (33 tumor types) (Li et al., 2013). The phospho-proteins measured include extensively annotated functional sites on kinases and transcription factors belonging to key signaling pathways implicated in cancers including Akt signaling, EGFR signaling, RAS-RAF pathway and hippo-signaling pathways. These sites can be used as a proxy for kinase signaling in cancer related pathways.

From the integration of large-scale pan-cancer genomic, transcriptomic and RPPA phosphoproteomic datasets we identified genes likely to drive changes in kinase signaling. These associations were found to be enriched in known kinase-substrate relationships and were then systematically filtered to select reliable associations that also replicated by analyzing cancer cell line data. A top-ranked predicted regulator (ARHGEF17) of hippo-signaling (YAP) was experimentally verified in two breast cancer cell lines and through (phospho-)proteomic analysis was shown to play a role in regulation of hippo-signaling and MAP Kinase pathway having an apparent downstream effect on cytoskeleton organization and immune effector processes. In parallel, we also found evidence of kinase addiction in cancer cell lines and identified inhibitors for targeting of ‘kinase-dependent’ cell lines. This work suggests that copy number status of genes may be indicative of kinase activity differences and predictive of sensitivity to kinase inhibition. In the long-term this strategy can be used to stratify patients for targeted therapy based on the copy number status of driver genes.

## Results

### Copy number alterations associated with phosphorylation changes in tumors

We have developed a computational method to predict genes driving changes in phosphorylation signaling in patient derived cancer samples (**Figure 1a**). A genetic association model was built to predict phosphorylation changes in tumor samples, obtained from RPPA data (TCPA), using CNVs as predictors of signaling events. The changes in phosphorylation, as measured in RPPA, could be either due to changes in total protein or changes in the stoichiometry of phosphorylation. In order to focus on changes in stoichiometry we restricted our analysis to 37 phosphosites which also had matched protein abundances in the RPPA Datasets for normalization purposes (**Methods**). CNVs and gene expression levels were obtained for 15,524 protein-coding genes after excluding genes whose CNVs have been predicted to be post-transcriptionally attenuated (Gonçalves et al., 2017). We then associated CNVs with phosphosite activity of 37 phosphosites across 5,598 tumor samples from 17 tumor types, taking into account total protein abundance and tumor type as covariates (**Methods**). Since copy number aberrations cover large chromosomal regions, phosphosite changes will show significant CNV associations with many genes in each co-amplification region, many of which are likely spurious. As an example, the genes found to be associated with changes in phosphorylation of Akt1 are shown in **Figure 1b**. On average, each phosphosite was predicted to be associated with 419 genes (10% FDR, Effect size > 0.05).

**Figure 1.**
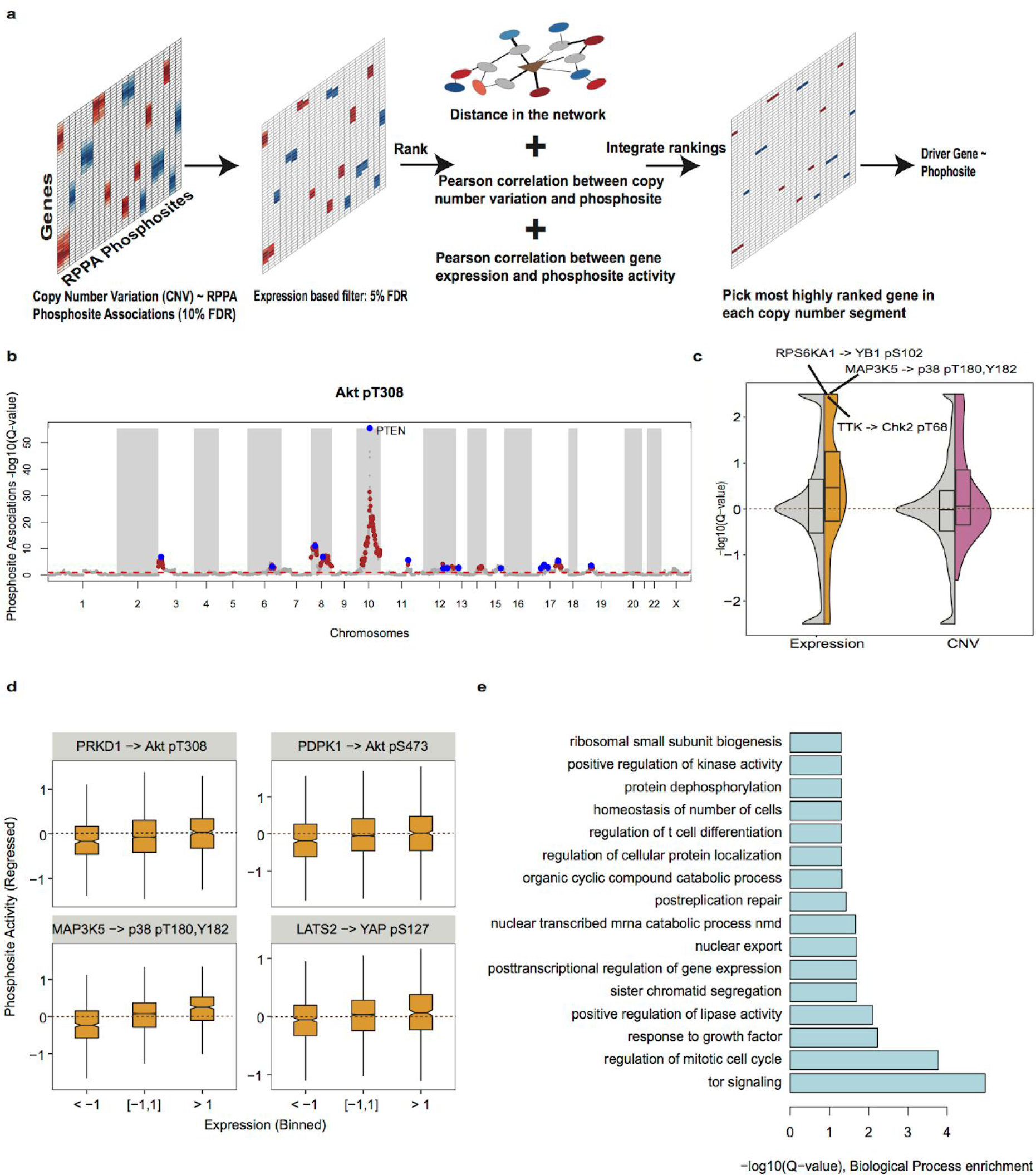
Prediction of regulators of phosphosite activity based on somatic DNA copy number aberrations and gene expression. (a) Key steps in the gene (CNV/Expression)~phosphosite phosphorylation association method. Red colour indicates strength of positive associations while blue colour represents strength of negative associations. Edge thickness are scaled by weights in the schematic network. (b) A Manhattan plot showing the genome-wide associations between gene copy number changes and Akt phosphosite levels (Akt pT308). Dashed red line indicates cutoff for significant copy number based associations (10% FDR). Genes highlighted in brown are supported by gene expression based associations. Genes highlighted in blue are the top-ranked gene within each genomic region. (c) significance of correlation between CNV (in purple) or mRNA (in yellow) levels with phosphosite changes for known kinase-substrate interactions (coloured) or any gene-phosphosite pair (in grey). (d) RPPA phosphosite levels, after regressing away protein changes, binned by mRNA levels of regulators in pan-cancer data from TCGA/TCPA for known regulator~substrate relationships. (e) Gene ontology over-representation analysis of predicted regulators.

Before filtering for spurious associations due to co-amplification, we tested if the CNV~phosphosite associations can recapitulate prior knowledge on kinase-substrate relationships. For this we analysed 118 known kinase-substrate relationships curated in the PhosphositePlus (Hornbeck et al., 2012) database involving the 37 phosphosites analysed here. Overall, we observe that known kinase~phosphosite pairs tend to have significant association more often than random expectation (P-value < 0.01) using both CNV and expression levels (**Figure 1c**). For instance the phosphorylation of YAP pS127, after regression for YAP total protein levels, showed a significant correlation with both expression and copy number changes of LATS2 kinase (Meng et al., 2016). Similarly, expression of PRKD1 (Scheid et al., 2002), PDPK1 (Sato et al., 2002), and MAP3K5 (Gu et al., 2009) kinases are positively correlated with Akt T308, Akt pS473 and p38 pT180,Y182 phosphosites respectively (**Figure 1d**).

Given that copy number alterations affect multiple genes within altered regions, many of the above defined associations are likely to be spurious passenger associations. We then sought to identify the most likely ‘driver’ gene controlling the downstream kinase signaling change for each region. Firstly, we selected CNV associated genes whose expression was also associated with the phosphosite activities (FDR < 5%). We then selected the top ranked associated genes within each chromosome region based on a combined rank from 3 measurements: 1) the CNV and 2) gene expression association effect size and 3) the gene~phosphoprotein distance in a functional interaction network (**Methods**). We exemplify in **Figure 1b** the genome-wide gene associations found as putative regulators of AKT1 pT308 phosphorylation levels and the selected driver genes predicted within each segment. For this site we recover the very well-known PTEN regulator(Lin et al., 2013), along with other novel candidate regulators. Using this approach an average of 8 predicted ‘driver’ genes per phosphosite were obtained, with each ‘driver’ gene as a representative of a genomic segment. All phosphosites except for PEA15 pS116 had at least one gene as a predicted regulator and 12 phosphosites including those on CHEK1, CHEK2, AKT, MAPK and YAP were predicted to be regulated by more than 10 driver genes.

We obtained a total of 303 associations between driver genes and regulated phosphosites in the pan-tumor dataset (**Supplementary Figure S1 and Supplementary Table S1**). These associations comprised of 264 driver genes among which 11% (29/264) of driver genes had more than one association. The driver genes showed significant enrichment for genes involved in *cell cycle process*, *TOR signaling*, *regulation of protein kinase activity* and *positive regulation of response to DNA Damage stimulus* (Fisher Adjust *p*-value < 0.01; **Figure 1e**). The driver genes were also enriched for kinases (19/264; Fisher Adjust *p*-value < 0.01) including FLT1, PTK6 and YES1 and phosphatases (11/264; Fisher Adjust *p*-value < 0.01) including PPA2 and PPP1CC. Consistent with their broad spectrum activity, phosphatases were found associated with multiple phosphosites when compared to other gene types (Wilcoxon rank-sum *p*-value < 0.05). For instance, PPP1R3B was predicted to regulate several phosphosites including p90RSK, Tuberin and FOXO3 (**Supplementary Table S1**). Similarly PTEN was predicted to regulate both Akt phosphosites (S473 and T308) and GSK3AB pS21,9. We also found a significant enrichment for known cancer driver genes from COSMIC database in our predicted driver genes list such as CDK4, EGFR, PTK6 and CCND1 (29/264; Fisher Adjust *p*-value < 0.01).

Several recurrent protein missense genetic variants are well known to cause changes in kinase signaling (Creixell et al., 2015). Given that some of the copy-number changes can co-occur with missense variants, we tested whether any of the 303 CNV associations with phosphosites can be explained by missense variants. From 1002 genes having missense variants in at least 50 patient pan-cancer samples we found 24 associations between recurrently mutated genes and phosphosite levels (**Supplementary Figure S2**). These include a positive association between EGFR mutation and EGFR pY1068 and HER2 pY1248 activity. Similarly, missense mutations in NRAS were positively associated with MAPK pT202, Y204 and mutations in KRAS were positively associated with MEK1 pS217, S221. We then asked if any of the CNV associations can be fully explained by one of the recurrently mutated genes. While in some cases there was a decrease in significance for the CNV~phosphosite associations, in only 6 cases the CNVs were not a significant added predictor. For example, the association between PDPK1 CNV with mTOR pS2448 activity could be explained by missense mutations in IL7R. Similarly, association between ZMYM1 CNV and EGFR pY1068 could be explained by mutations in EGFR.

### Replication of predicted drivers of signaling changes in cancer cell lines

To investigate the reliability of the predicted associations from the pan-tumor analysis, we attempted to replicate the significant associations in the cancer cell line cohort from CCLE with publicly available copy number (Barretina et al., 2012), expression (Barretina et al., 2012) and phosphorylation (Zhao et al., 2017) information. We interrogated two independent RPPA datasets from DepMap and MCLP to test the replicability of these associations in cancer cell lines. A total of 130 out of 303 of the predicted driver gene ~ phosphosite were replicated in the larger set of 890 cancer cell lines in DepMap (10% FDR; **Supplementary Table S1**). Separately, 66 out of 303 predicted driver gene ~ phosphosite were replicated in the smaller set of 319 cancer cell lines in MCLP (10% FDR; **Supplementary Table S1**), using the mRNA levels of the predicted driver genes. Differences in predictions may be partially due differences in sample size and/or differences in heterogeneity of the tumor versus cell line samples, such as presence of non-tumor cells or changes during in vitro culturing. To assess the overall predictive power of activity of individual phosphosite, we trained models of phosphosite activities as a linear combination of expression profiles of the predicted driver genes (**Methods**). For 8 out of 37 phosphosites, at least 20% of the variance in phosphosite activities in DepMap cell lines could be significantly explained by the expression of predicted driver genes in the cell lines. One of these phosphosite, MEK1 pS217,S221, at least 20% of variance in activity in DepMap cell lines could be predicted using linear model trained on the TCGA tumor dataset. For 12 out 37 of phosphosites, at least 20% of the variance in phosphosite activities in MCLP cell lines could be significantly explained by the expression of predicted driver genes (**Figure 2a**). For three of these phosphosites, HER3 pY1289, EGFR pY1068 and YAP pS127, at least 20% of variance in activity in MCLP cell lines could be predicted using linear model trained on the TCGA tumor dataset (**Figure 2a**).

**Figure 2.**
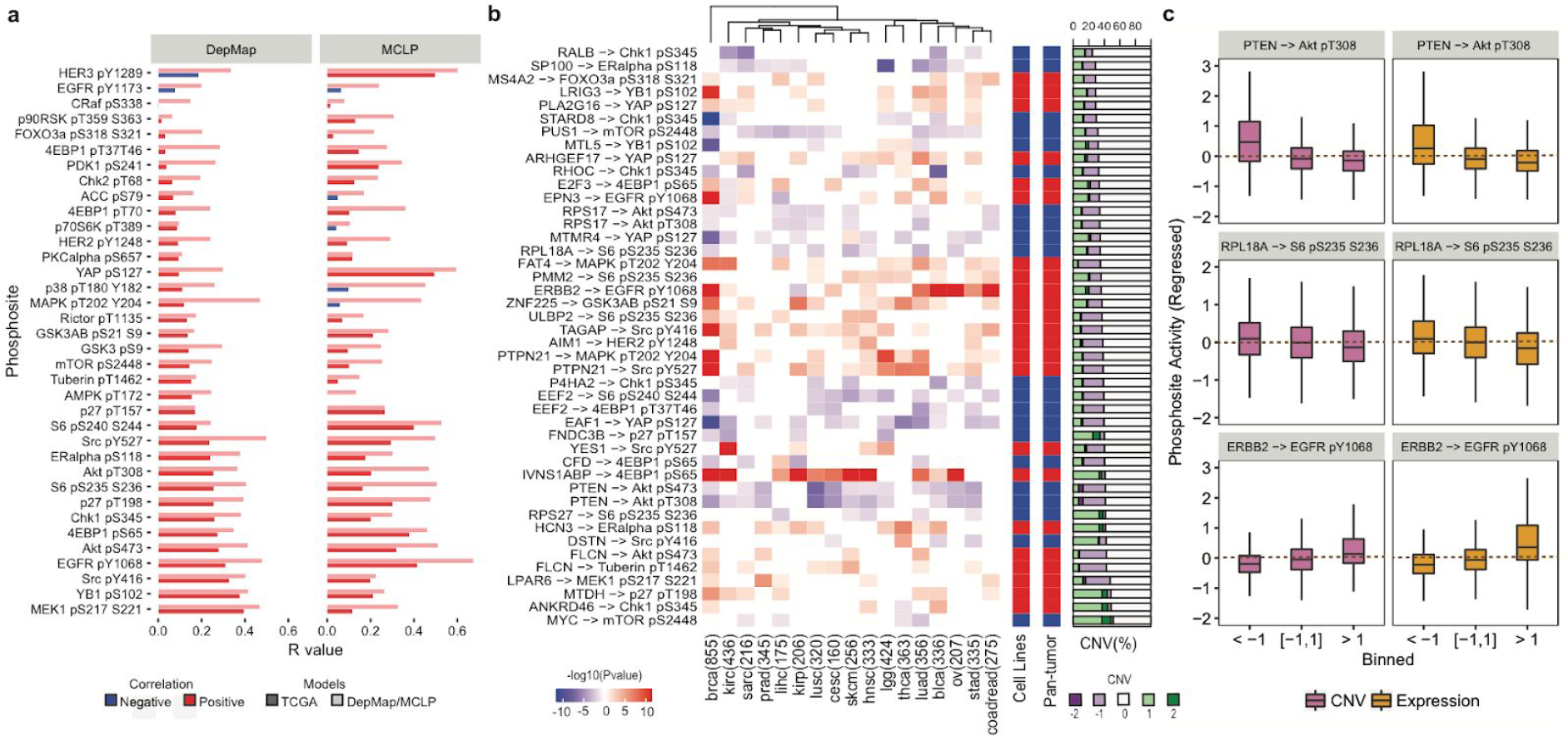
Shortlisted gene~phosphosite pairs replicated in cancer cell line data. (a) Variation explained in phosphosite levels as predicted by a linear model for each phosphosite based on expression of putative driver genes. Linear models derived from TCGA (dark), DepMap (light) and MCLP (light) based RPPA data were used to predict phosphosite activity on cell line based expression data and correlated with the measured DepMap and MCLP RPPA measurements. (b) Shortlist of gene~phosphosite pairs predicted in pan-tumor analysis and replicated in both DepMap and MCLP cell lines. (c) Changes in phosphosite levels (after regressing out protein changes), binned by mRNA levels or copy number levels of predicted regulators for some top ranked and replicated pairs.

A stringent set of 44 driver gene~phosphosite pairs were defined which were replicated in at least 20% of different tumor types (4/18) and also replicated independently in two different cancer cell line datasets (**Figure 2b**). Among the replicated associations were well-known examples (**Figure 2b and 2c**) such as loss of PTEN correlated with increase in phosphorylation of Akt pT308 and Akt pS473 (Lin et al., 2013) and the EGFR amplification correlated with HER2 pY1248 phosphosite activity(Dittmar et al., 2002). In addition to these known examples, we were able to identify several novel potential regulators of phosphosite levels which have been previously reported to play a key role in cancer development and/or progression. For instance, MTDH gene known to be involved in regulation of several signaling pathways including Akt, Wnt and MAPK leading to tumor metastasis(Emdad et al., 2013), was predicted to positively regulate p27 pT198 (cyclin-dependent kinase inhibitor) activity.

### Evidence of kinase addiction in cancer cells

Having shown that copy number alterations lead to kinase signaling changes we next sought to find cases where such signalling differences create potential genetic vulnerabilities. Previous studies have proposed that cancer cells have a tendency to become dependent on the activity of proteins such as kinases (Tyner et al., 2013) and are therefore likely to be sensitive to loss of the kinase gene. We exploited the recently generated RNAi (Tsherniak et al., 2017) and CRISPR (Meyers et al., 2017) knock-down and KO data for cancer cell lines in Project Achilles to test the generality of this concept. RPPA phosphosites were classified into activating (25 sites) and inhibitory (9 sites) based on evidence in literature (Hornbeck et al., 2012; Korkut et al., 2015). Protein activities as measured by phosphosites in DepMap were then correlated with sensitivity to loss of the corresponding gene across 406 cancer cell lines in the CRISPR screen. Increased protein activity in a given cell line showed an overall tendency for increased essentiality of the gene in the respective cell line (median log10 P-value = −0.44; **Figure 3a**). Activation sites showed an overall tendency for negative correlation and inhibitory sites show an overall tendency for positive correlation (**Figure 3b**). The protein activity dependencies varied across different genes and were recapitulated for several proteins using a RNAi data with matched RPPA phosphoproteomic measurements (**Figure 3b**). The strongest reproduced activity dependencies were observed for YB1, AKT, Her2, ER alpha and MEK with other strong dependencies observed in at least one study. We repeated the same analysis of associations between RNAi/CRISPR screens with RPPA data from MCLP dataset and observed the same trend of phosphosite activity to be correlated with cellular sensitivity to loss of phosphosite gene although stronger associations were observed for different phosphosites such as YAP pS127 (**Supplementary Figure S3**). This analysis confirms that, on average, cancer cell lines show increased dependency on the activity of kinases and other proteins and are sensitive to their inhibition.

**Figure 3.**
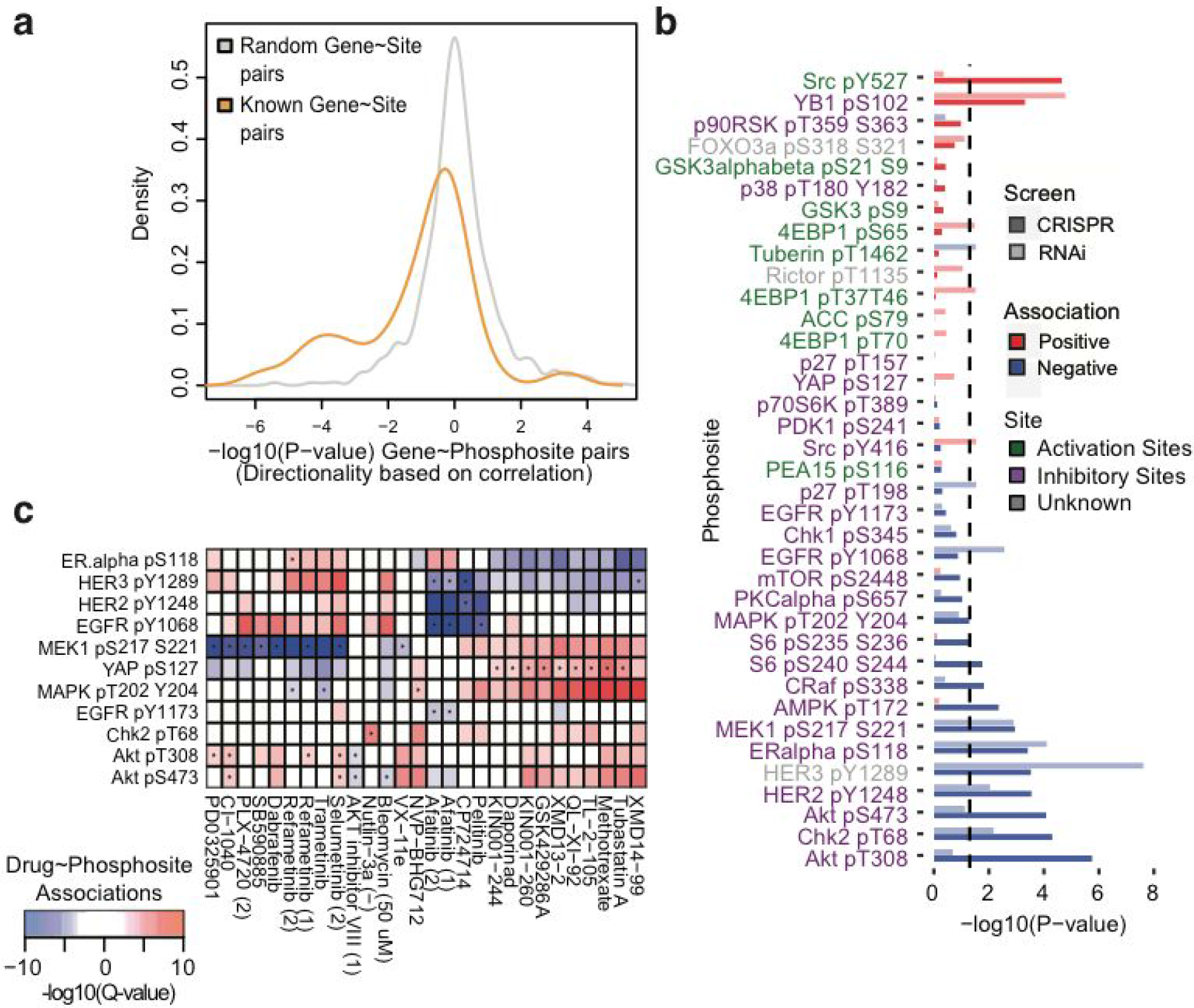
Evidence of kinase addiction in DepMap cancer cell lines. (a) Distribution for the significance of association (−log_10_(p-value)) between phosphosite activity and gene essentiality (Crispr). (b) Significance of association (−log10(p-value)) between phosphosite activity and gene essentiality for each phosphosite using both RNAi and CRISPR Screen. (c) Association between phosphosite activity and drug sensitivity (1% FDR). Positive and negative associations and also supported by RNAi~Drug response are indicated in red and blue respectively. * indicates associations also supported by CRISPR~Drug association.

We next investigated drug response datasets to interrogate the relationship between phosphosite levels and drugs. We correlated protein activity as measured by 37 phosphosites with sensitivity of 265 Drugs in 746 cancer cell lines from GDSC database (Yang et al., 2013), with 30 phosphosites showing significant correlation with sensitivity of at least one drug target (**Supplementary Figure S4**, FDR < 5%). A substantial fraction of these phosphosites (18/30) were associated with sensitivity to more than 25 drugs. For instance, cancer cell lines with high MEK1 pS217,S221 activity were more sensitive to the majority of RAF/MEK/ERK inhibitors (such as Refametinib, SB590885). Similarly, YAP pS127 activity correlated with Methotrexate inhibition. We further refined these associations to identify 870 cases where the drug sensitivity was also correlated with sensitivity to loss of the kinase/phosphosite gene in either RNAi or CRISPR screen (**Figure 3c**). In 31 of these cases there was a concordance in both gene perturbation screens. For instance, cell lines sensitive to EGFR inhibitors (Pelitinib and Afatinib), as predicted by high EGFR phosphosite activity, were also sensitive to loss of EGFR gene in both CRISPR and RNAi datasets. Similarly, cell lines sensitive to ERBB2 inhibitor (CP724714) also correlated with high HER2 pY1248 and HER3 pY1289 phosphosite activity, were also sensitive to loss of ERBB2 gene in both CRISPR and RNAi datasets. Many of these associations between phosphosite activity and drug response were also replicated using the RPPA data from MCLP (**Supplementary Figure S3**).

### CNVs as predictors of kinase related genetic susceptibility

Our results indicate that copy number changes in genes can be predictive of changes in phosphosite levels and that cancer cells can become dependent on activity levels as measured by these phosphosites. Therefore, copy number changes correlated with kinase activity should also be predictive of kinase susceptibility. Indeed, we found several examples where the CNVs predictive of protein activity differences also correlated with the down-regulation, knock-out or inhibition of the corresponding protein. For instance, ERBB2 and EPN3 CNVs were predictive of EGFR pS1068 phosphosite activity and also correlated with sensitivity to loss of EGFR gene and sensitivity to EGFR inhibitors including Afatinib and Gefitinib (**Figure 4a and Figure 4b**).

**Figure 4.**
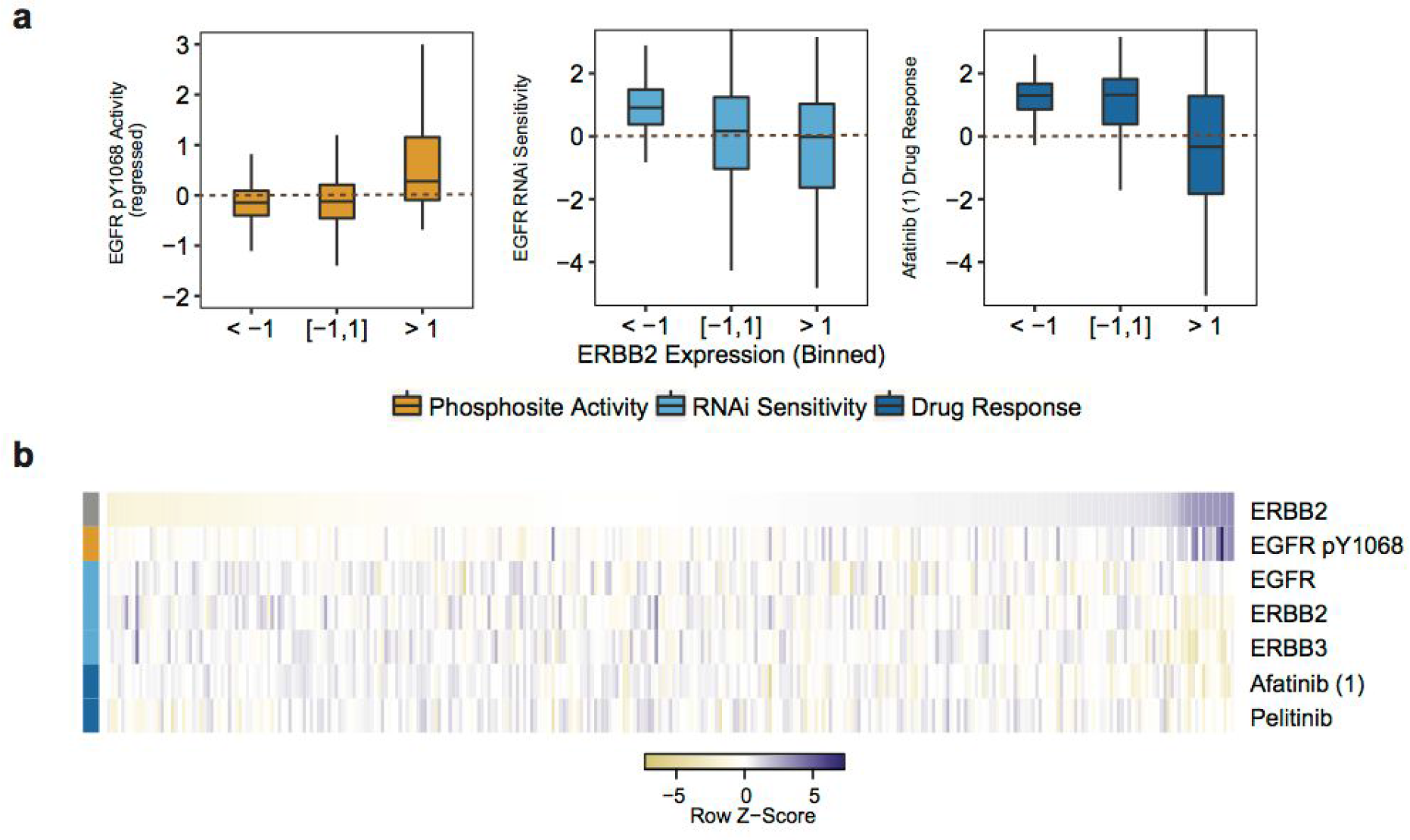
CNVs as predictors for kinase related genetic susceptibility. Levels of EGFR pY1068 activity, RNAi sensitivity to loss of EGFR (and ERBB2 and ERBB3) gene and sensitivity to related kinase inhibitors when (a) binned or (b) ordered according to ERBB2 expression levels.

From the CNV~phosphosite association analysis, we have identified 12 genes as potential drivers of YAP pS127 phosphorylation, 5 of which were replicated in cancer cell lines (**Figure 5a**). We ranked the genes based on their association with YAP phosphosite activity in a multiple linear regression model using tumor (TCGA) and the two different cell line RPPA datasets (**Figure 5b**). In the combined model, the CNVs of ARHGEF17 and PLA2G16, showed the strongest association of YAP pS127 activity in tumor and at least one cell line dataset (**Figure 5b**). Expression of ARHGEF17 and PLA2G16 was also associated with sensitivity to loss of YAP gene and other core components of hippo-signaling pathway such as LATS2 (**Figure 5c,d,e**). YAP, a transcription co-factor, is a key component of the hippo-signaling pathway (Gumbiner and Kim, 2014). Phosphorylation of YAP at S127 leads to the retention of YAP in the cytoplasm (Zhao et al., 2007) and further phosphorylation of YAP at S381 and S384 is associated with YAP degradation (Zhao et al., 2010). For these reasons, the phosphorylation of S127 should result in a decrease in the transcriptional activity of YAP. However, phosphorylation at S128 can also block the 14-3-3 mediated cytoplasmic retention of S127 phospho YAP (Moon et al., 2017). In order to disambiguate the transcriptional activity changes as detected by the S127 RPPA antibody we studied the gene expression changes of 45 positively regulated targets of YAP (Kim et al., 2015). We observed on average a positive correlation between the changes in expression of these genes and the changes in total YAP, total phosphorylated YAP and also the changes in phosphorylated YAP after normalizing for YAP protein levels (**Supplementary Figure S5**). This was the case in both the patient tumor samples and MCLP cell lines (**Supplementary Figure S5**). This would suggest that the RPPA antibody for S127 is positively associated with YAP activity. In line with this there is a positive correlation between the RPPA YAP S127 levels and the sensitivity of cells to knock-down or knock-out of YAP.

**Figure 5.**
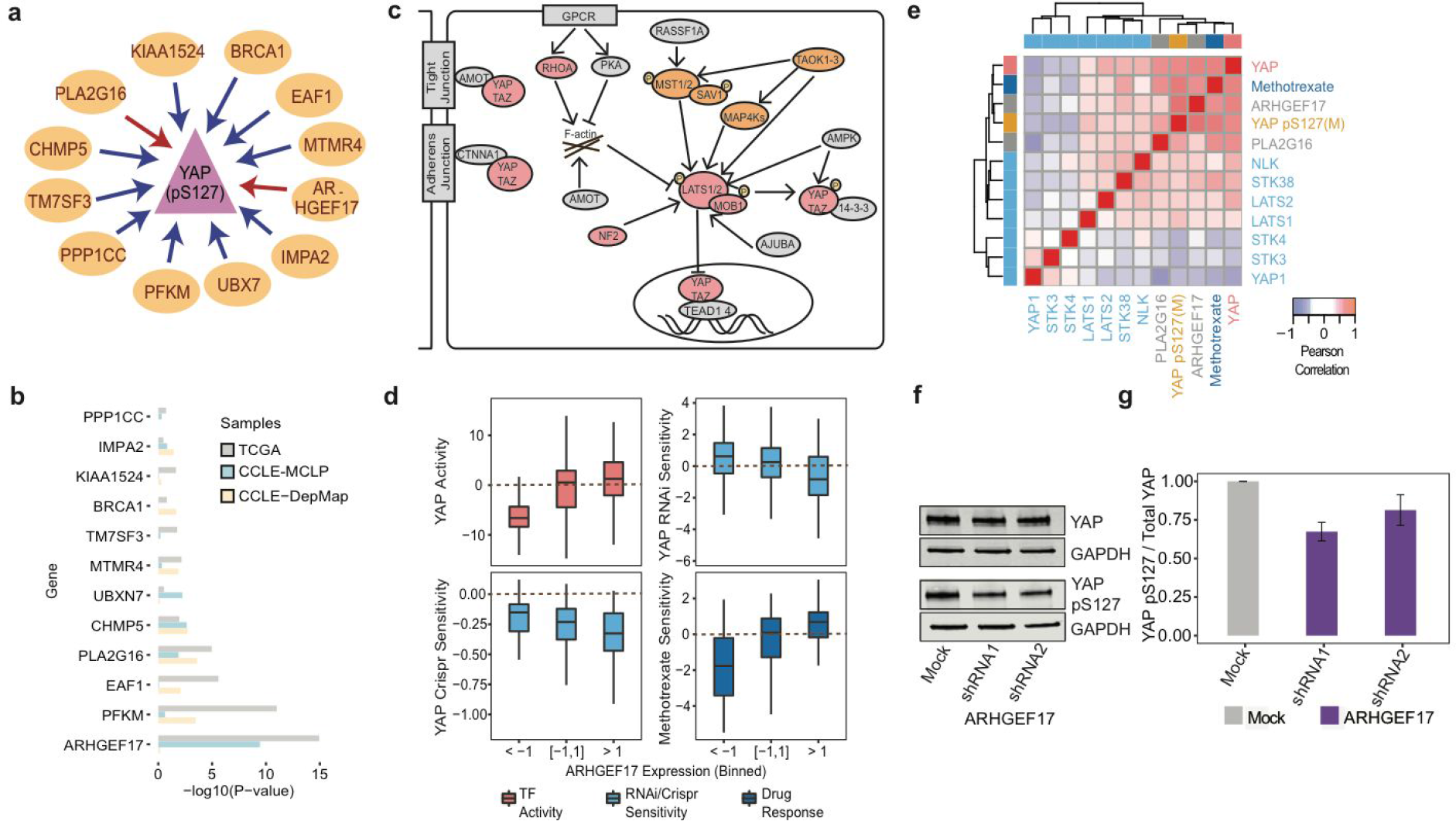
Experimental validation of predicted regulators of YAP pS127 in breast cancer cells. (a) Schematic representation of predicted regulators of YAP pS127. Red and blue arrows indicate positive correlation and negative correlation respectively. (b) Contribution of individual regulator to YAP pS127 activity in a multiple linear regression model. (c) Schematic representation of Hippo-signaling pathway adapted from Plouffe et al.(Plouffe et al., 2016) where red and orange circles indicate critical/important components of YAP/TAZ regulation. (d) Levels of YAP co-factor activity, RNAi/Crispr sensitivity to loss of YAP gene and sensitivity to predicted drugs when binned according to ARHGEF17 levels. (e) Pairwise correlation coefficient values between the: expression levels of PLA2G16 and ARHGEF17 (grey); sensitivity to drugs (dark blue); YAP transcriptional activity (red); phosphorylation levels from MCLP dataset (orange); and sensitivity to knockdown of hippo pathway members (light blue). (f) Western blot analysis of YAP protein and YAP pS127 levels and (g) quantification of relative YAP pS127 levels on knockdown of PLA2G16 and ARHGEF17.

We experimentally tested the effect of loss of ARHGEF17 and PLA2G16 genes on YAP pS127 phosphosite levels. We selected cell lines with different baseline levels of YAP phosphorylation levels and confirmed that YAP pS127 levels were low in MCF7 and MDA-MB-361 cells and high in MDA-MB-468 and T-47D cells consistent with the measurements in MCLP RPPA phosphoproteomics dataset from cancer cell line cohort (**Supplementary Figure S6**). Effective lentiviral shRNA mediated knockdown of both ARHGEF17 and PLA2G16 in MDA-MB-468 and T-47D cells was confirmed using qRT-PCR (**Figure 5f** and **Supplementary Figure S6**). Knockdown of ARHGEF17 resulted in a small but significant decrease in the levels of YAP pS127 in both cell lines which was consistent with the predictions (p < 0.05; **Figure 5g**). Loss of PLA2G16 had also a reproducible but modest effect on YAP pS127 (**Supplementary Figure S6**).

### ARHGEF17 knockdown disrupts YAP activity and down-regulates immuno-responsive pathways

To further dissect the regulatory role of ARHGEF17 in hippo-signaling we performed proteomics and phospho-proteomics of MDA-MB-468 and T47D cells treated with shRNA targeting ARHGEF17 along with their matched controls (**Figure 6a**). Differential phosphosite analysis data showed significant changes in 3182 and 259 phosphosites after ARHGEF17 knockdown (5% FDR) in MDA-MB-468 and T47D cells respectively (**Supplementary Table S2**). These include significant decrease in multiple YAP phosphosites in both cell lines (**Figure 6b**,**S6**). We next estimated the changes of kinase activity after ARHGEF17 knockdown by studying the changes in known kinase target phosphosites(Ochoa et al., 2016). The substrates of four kinases, NDR1, CK2A1, P38G and PKG showed significant down-regulation in both MDA-MB-468 and T47D cells on knockdown of ARHGEF17 (**Figure 6c**). NDR1 (STK38) is reported to be a kinase directly interacting with YAP pS127 (Zhang et al., 2015), and P38G, a ser/thr MAP kinase which has been previously associated with YAP signaling (Huang et al., 2016; Liu et al., 2016). Notably, this association between the two pathways is also observed in pan-cancer TCGA data with a strong correlation between YAP pS127 and p38 pT180,pY182 (P38A) (**Supplementary Figure S7**).

**Figure 6.**
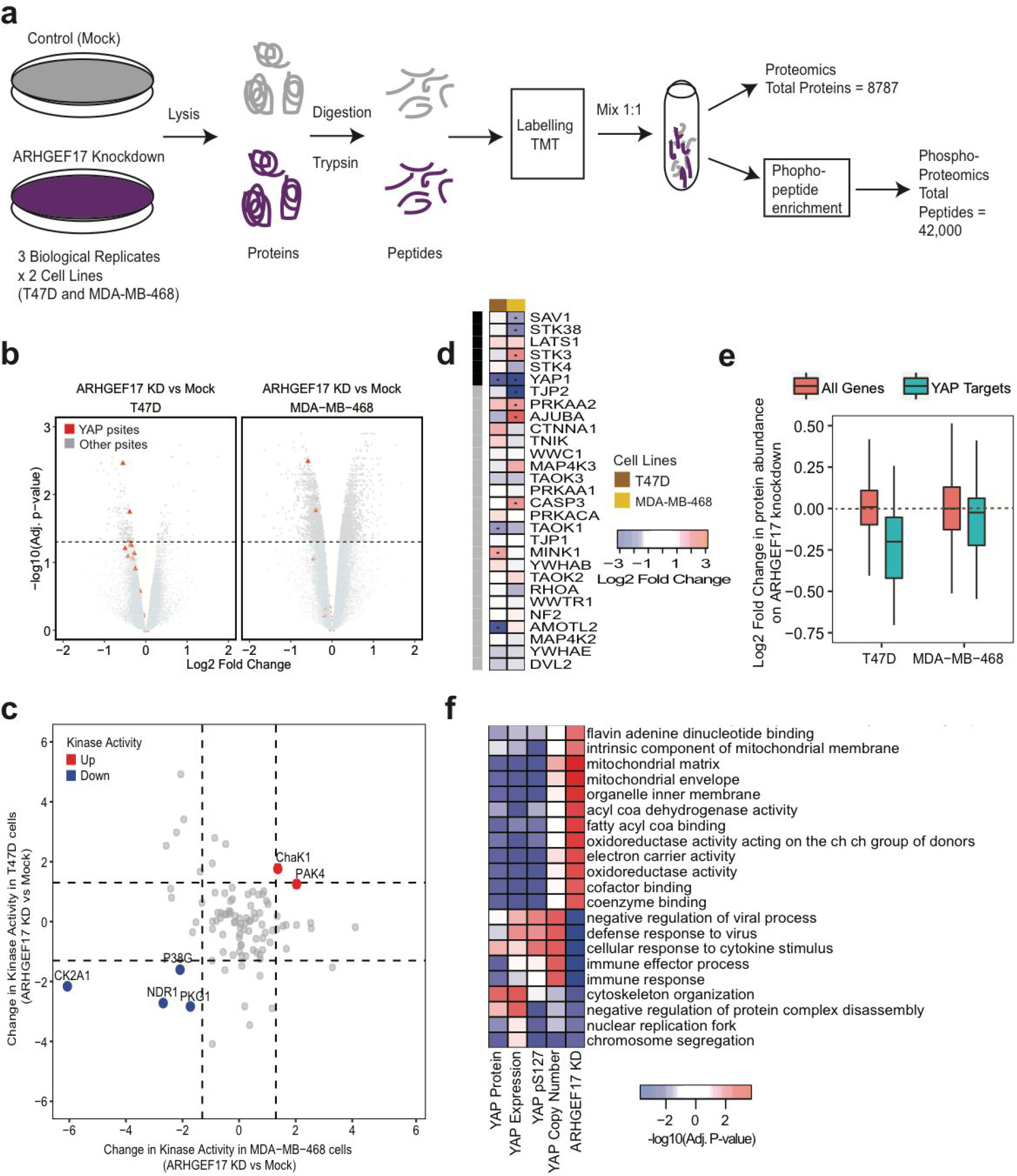
ARHGEF17 regulates hippo-signaling pathway. (a) Experimental design of (Phospho-)Proteomics data generation on knockdown of ARHGEF17. (b) Differential analysis of phosphorylation data in T47D and MDA-MB-468 cells with YAP phosphosites highlighted in red. (c) Change in predicted kinase activity after ARHGEF17 knockdown. (d) Changes in protein abundances of component of hippo-signaling pathway after ARHGEF17 knockdown. (e) Comparisons of YAP cofactor activity on ARHGEF17 knockdown estimated from the proteomics data. The downstream targets of YAP were obtained from Kim et al. (Kim et al., 2015) (f) Gene Set Enrichment Analysis of biological processes from gene expression changes after ARHGEF17 knockdown or genes whose expression levels correlates with different measurements of YAP activity or levels in the TCGA patient data.

We then studied the protein abundance changes and particularly that of components of hippo-signaling pathway upon ARHGEF17 knockdown. Differential protein abundance analysis showed significant changes in 1754 and 308 proteins after ARHGEF17 knockdown (5% FDR) in MDA-MB-468 and T47D cells respectively (**Supplementary Table S2**). ARHGEF17 protein levels was reliably decreased in both cell lines with 5-fold and 10-fold down-regulation in MDA-MB-468 and T47D cells respectively. We also observed a significant decrease in YAP protein levels in both breast cancer cell lines which may explain most of the changes in phosphorylation. Several “core” components of hippo pathway including STK3, STK38 and SAV1 were also found to be significantly dys-regulated in MDA-MB-468 cell lines (**Figure 6d**). Importantly we again accessed the changes in YAP co-factor activity, derived from positively regulated targets of YAP observing a strong decrease in T47D cells (P-value < 0.01) and a more modest decrease in MDA-MB-468 cells (**Figure 6e**). Finally, we analyzed the changes in protein abundance to identify biological processes that were dys-regulated upon ARHGEF17 knockdown. Gene set enrichment analysis of biological processes showed down-regulation of *cytoskeleton organization*, *defense response to virus* and *immune effector process* and up-regulation of *electron carrier activity* and *fatty acyl coA binding* pathways (**Figure 6f**). In order to relate these changes to the tumor samples we correlated the gene expression changes occurring for the same gene sets with YAP copy number, mRNA, protein and YAP pS127 levels. In line with the observations from the ARHGEF17 knockdown experiments YAP activity measurements were associated with changes in gene expression of the same biological processes in the patient tumor samples (**Figure 6f**).

Together these analyses indicates a role for ARHGEF17 in regulation of hippo-signaling pathway and hippo pathway affecting cytoskeleton organization and immune response pathways in cancer cells.

## Discussion

We propose here a genetic association approach to identify genes whose expression changes can drive differences in kinase-signaling. We note that such “regulators” are unlikely to be direct as they are found through genetic association for differences in steady state phosphorylation levels. We make use of copy number variation, instead of recurrent missense mutations, given that they form a critical mass of genomic aberrations encountered in multiple tumor types and compared to point mutations the impact of copy number changes on an individual gene is easier to interpret. For the majority of genes, their copy number profiles tend to be well correlated with gene expression data (Hyman et al., 2002). While the degree of changes and interpretability makes CNVs an attractive choice for association analysis, these changes typically involve a large number of genes and in some cases entire chromosomal segments. In these scenarios, it is a challenge to identify the key driver gene of downstream signaling changes among the co-amplified or co-deleted block of genes. Here we have defined an approach to prioritize copy number events that are regulators of signaling pathways, showing that the method was able to recover previously known regulatory relationships. However, it is possible that the top-selected gene will not be the driver gene for the associated genomic region and we provide in the **Supplementary Table 1** the rank of genes associated with a phosphosite for each segment.

We selected for further analysis ARHGEF17 as a putative novel regulator of hippo-signaling pathway. ARHGEF17 is a Rho GTPase with a functional role in maintenance of actin cytoskeleton organization and focal adhesion (Mitin et al., 2013). In the tumor patient samples, cancer cell lines and in the direct knockdown experiments the levels of ARHGEF17 were positively correlated with the transcriptional activity of YAP. However, the relationship between YAP phosphorylation and protein levels is not easy to disentangle and we cannot rule out, based on the current set of results, that the ARHGEF17 mediated regulation of YAP is primarily driven by protein level changes.

The current study relied on antibody measurement of phosphorylation and protein abundance levels. More recently, mass spectrometry has started to be used to profile tumor samples in large scale for an increasing number of samples (Edwards et al., 2015). While the current limited set of samples makes it difficult to perform genome-wide association level analysis, as these efforts progress the approach described here can be applied to those data providing potentially with a much higher resolution description of how different CNVs rewire kinase signaling in cancer.

Cancer cells often become dependent on changes in the genes that led to the cancer development, a phenomena dubbed oncogene addiction. When a signaling pathway is observed to be hyperactive in a tumor, relative to healthy tissue, it is often assumed that such activation is a driving event and a requirement for cell viability. For this reason, cellular dependency on kinase activity has been exploited in many targeted therapies (Bhullar et al., 2018; Gross et al., 2015). We tested here the generality of such dependencies finding that on average it is observed although it is highly variable depending on the specific signaling protein. Our results suggest that the copy number profile of a cancer cell can, in principle, be used as a ‘molecular fingerprint’ to stratify those tumors more likely to be sensitive to specific kinase inhibition. Since copy number events do not occur in isolation, additional work will be needed to understand how the combinatorial effect of multiple mutations can change the way signaling network works.

## Methods

### Compilation of molecular and phenotypic data

Normalized copy number (log2) datasets (10,654 samples) and RNAseq expression datasets (9,548 samples) for 31 different tumor types generated as part of TCGA (Cancer Genome Atlas Research Network et al., 2013) consortium were obtained from cBioPortal(Gao et al., 2013) database. Normalized copy number (log2) dataset (995 cell lines) and RNAseq expression dataset (967 cell lines) were obtained from cancer cell lines (CCLE) (Barretina et al., 2012). Normalized RPPA (phospho)proteomics datasets for 31 different tumor types comprising of 7694 samples were obtained from TCPA (Li et al., 2013) database. Normalized RPPA (phospho)proteomics dataset were download for 651 cancer cell lines from MCLP (v1.1) (Li et al., 2017) and 899 cancer cell lines from DepMap.

Genome-wide RNAi (Tsherniak et al., 2017) screen and CRISPR (Meyers et al., 2017) screens were obtained from the Broad Institute portal (https://portals.broadinstitute.org/achilles). These datasets were generated by Project Achilles to catalog gene essentiality across cancer cell lines. The RNAi screen (Ach 2.20.2) has high coverage targeting 17,098 genes in 501 cell lines. Out of these, and 406 cell lines had DepMap-RPPA data and 166 cell lines had available MCLP-RPPA data (Li et al., 2017) permitting pan-cancer correlative analysis. The CRISPR screen (Avana CRISPR-Cas9) targets 17,670 genes across 517 cell lines. 420 of these cell lines also had DepMap-RPPA data and 163 cell lines also had MCLP-RPPA data. IC50 values of sensitivity to inhibitors/drugs were obtained from Genomics of Drug Sensitivity in Cancer Database (Yang et al., 2013). The dataset comprises of sensitivity values for 265 drugs across 746 cell lines. These cell lines were mapped with those in CCLE to identify 596 cell lines with DepMap-RPPA estimates and 259 cell lines with MCLP-RPPA estimates.

### Modelling the genetic association between gene copy number and phosphosite levels

A total of 6,558 samples were compiled with information on Copy Number, Expression and phosphorylation levels measured with RPPA for patient samples from TCGA. We restricted our analysis to 5598 samples from 17 tumor types which had more than 150 samples. We predicted associations between 18,350 CNVs and 37 RPPA phosphosites across 17 different tumor types using linear mixed effect models. Each linear model had phosphosite activity as response variable, CNV as the explanatory variable and matched protein abundance (same as phosphosite) and tumor type as covariates. A separate covariate model was run for each phosphosite. Two models were compared using a likelihood ratio test to predict the effect of a given CNV on phosphosite activity after taking into account the matched protein abundance and tumor specificity. A network-based score to rank genes was derived by calculating a network-based score between the CNV gene and phosphosite gene in the STRING network.

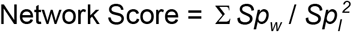

Where *Sp*_*w*_ and *Sp*_*l*_ are edge weights and length of the shortest path between the CNV gene and the phosphosite gene in the STRING network respectively.

Significant associations predicted in tumor samples were also tested in 890 cancer cell lines with RPPA and expression data in DepMap database and 319 cancer cell lines with Expression and RPPA datasets. To evaluate the quality of our predictions from the pan-tumor analysis, we predicted the phosphosite activity from expression data in cancer cell lines using models trained using TCGA, MCLP and DepMap data and correlated with the ‘true’ phosphosite activities measured independently in MCLP or DepMap databases.

### Cells lines and maintenance and RNA Interference

MDA-MB468, MDA-MB-361, T-47D and MCF7 were all maintained at 37°C with 5% CO2. All cells were grown in DMEM with 10% FBS. For serum starvation, cells were incubated in DMEM without other supplements for 24 h. ShRNA hairpins were designed using (http://sherwood.cshl.edu:8080/sherwood/) and cloned into the expression vector pZIP-SSFV-Ultra-Puro_ZsGreen (a generous gift of the Hannon Lab, CRUK CI). ShRNA plasmids together with pMDL, pCMV-Rev, and pVSV-G were used to produce virus in 293FT cells. Resulting viruses were used to infect and generate stable cell lines for both MDA-MB-468 and T-47D using puromycin selection. For shRNA primer details see supplementary methods

### Cell lysates, Immunoblotting and qRT-PCR Gene Expression Analysis

Cells were lysed in 50 mM Tris-Cl, pH 7.6; 150 mM NaCl, 1% NP-40 supplemented with Protease inhibitors (cOmplete EDTA free and PhosStop phosphatase inhibitor, Sigma). Cells lysates were centrifuged for 10 min at 4oC to remove insoluble debris. Lysates were separated using 4-20% Gradient gels (Miniprotein TGXTM Gel, Biorad) and immunoblotted using standard protocol. Primary antibodies used were YAP (4912S/Cell Signaling Technology, 1:1000), p-YAP (S127)(4911S/ Cell Signaling Technology, 1:1000) and GAPDH (MAB374/EMD Millipore, 1:5000). Blots were probed with mouse or rabbit specific IRDye 800 (LiCOR) and acquired, and quantified, using Odyssey Classic. Total RNA was isolated and purified from cells using miRNeasy Mini Kit (Qiagen). cDNA was synthesised using the High Capacity RNA-to-cDNA kit (ABI) according to manufacturer’s instructions. qRT-PCR was performed using Fast SYBR™ Green Master Mix (ABI) on the QuantStudio 12K Flex Real-Time PCR System (ABI). Relative expression levels were defined using the DDCt method and normalising to 36B4/RPLP0. For PCR primers details see supplementary methods.

### Proteomics and Phosphoproteomics analysis

Samples were collected for 3 biological replicates of ARHGEF17 knockdown and after protein sample preparation, proteins were trypsin-digested and TMT labelled. Each sample was split for full proteome analysis (10%) and phosphoproteome analysis (90%). The portion for the phospho-proteome analysis was fractionated using basic reversed phase C18 chromatography, each fraction was enriched using the High-Select Fe-NTA Phosphopeptide Enrichment kit (Thermo Scientific, #A32992) according to manufacturer’s instructions. The Phosphopeptide-enriched fractions were analysed on the Q-Exactive HF (Thermo Scientific) The total proteome was fractionated using basic reversed-phase C18 chromatography and analysed with the Fusion-Lumos Orbitrap mass spectrometer (Thermo Scientific). Both systems were configured with a Dionex Ultimate 3000 RSLC nanoHPLC system. Both total proteome and phospho-proteome raw files were processed with the SequestHT search engine on Proteome Discoverer 2.1 software for peptide and protein identification.

Quantification of 65,535 peptides were generated for both t47d and mb468 cells. The data were quantile normalized separately for each cell line and summarized at protein level based on the median abundance of all the peptides mapped to each protein in a sample. Therefore, a total of protein abundance measurements were obtained for 7,963 and 7,943 proteins from T47D and MDA-MB-468 cells respectively with 7,119 proteins common between the two cell lines. Gene set enrichment analysis (GSEA) (Barbie et al., 2009) to identify to differentially regulated biological processes in proteome data was performed using clusterProfiler (Yu et al., 2012) package in R. Annotations for biological processes were obtained from Molecular Signature Database (MSigDB) (Liberzon et al., 2011). Downstream targets of YAP transcription co-factor were obtained from Kim et al (Kim et al., 2015) from which we retained only positively regulated targets for our analysis.

The phosphoproteomics measurements were obtained for 41,249 and 40,754 phosphopeptides in t47d and mb468 cells respectively. These were quantile normalized and differential phosphoproteome analysis was performed using the limma (Ritchie et al., 2015) package in R. Activity of kinases were estimated using the changes in phosphorylation of known substrates on the phosphoproteomics experiments. Known kinase-substrate relationship in literature have been collated from the PhosphositePlus (Hornbeck et al., 2012) database and only kinases with at least five known substrates with measurements in the phosphoproteomics dataset were analysed. There are several statistical variants of Kinase Set Enrichment Analysis (KSEA) approach and we used the KS statistics which is faster and has been shown to perform comparably with traditional ssGSEA based statistic (Hernandez-Armenta et al., 2017).

## Acknowledgments

We acknowledge Dr. Matthew Garnett (Wellcome Trust Sanger Institute), Dr. Emmanuel Goncalves (Wellcome Trust Sanger Institute) and Dr. Colm Ryan (University College Dublin) for their insightful feedback on the manuscript. D.M. is a fellow of the joint EMBL-EBI & NIHR Cambridge Biomedical Research Centre Postdoctoral (EBPOD) program.

## Data Access

All proteomics and phosphoproteomics datasets generated as part of this study are deposited in PRIDE database (Accession No:).

